# Characterization of the flagellar collar reveals structural plasticity essential for spirochete motility

**DOI:** 10.1101/2021.08.23.457452

**Authors:** Yunjie Chang, Hui Xu, Md A. Motaleb, Jun Liu

## Abstract

Spirochetes are a remarkable group of bacteria with distinct morphology and periplasmic flagella that enable motility in viscous environments, such as host connective tissues. The collar, a spirochete-specific complex of the periplasmic flagellum, is required for the unique spirochete motility, yet it has not been clear how the collar assembles and enables spirochetes to transit between complex host environments. Here, we characterize the collar complex in the Lyme disease spirochete *Borrelia burgdorferi*. We discover as well as delineate the distinct functions of two novel collar proteins, FlcB and FlcC, by combining subtractive bioinformatic, genetic, and cryo-electron tomography approaches. Our high-resolution *in-situ* structures reveal that the multi-protein collar has a remarkable structural plasticity essential not only for assembly of flagellar motors in the highly curved membrane of spirochetes but also for generation of the high torque necessary for spirochete motility.

## Introduction

Spirochetes are a group of bacteria that cause several serious human diseases, such as Lyme disease (*Borrelia burgdorferi*), syphilis (*Treponema pallidum*), periodontal disease (*Treponema denticola*), and leptospirosis (*Leptospira interrogans*). Spirochetes have a distinctive spiral or flat-wave morphology (Charon et al., 2012; Nakamura, 2020). Enclosing the cell is a multilayered envelope including the outer membrane, peptidoglycan layer, and cytoplasmic membrane. The motility of spirochetes is unique among bacteria, as the whole cell body rotates without any external apparatus. Furthermore, this motility is crucial for host tissue penetration, virulence, and transmission of spirochetes (Lambert et al., 2012; Li et al., 2010; Lux et al., 2001; Motaleb et al., 2015; Sultan et al., 2015). The main organelle responsible for spirochete motility is the periplasmic flagellum, which rotates between the outer membrane and peptidoglycan layer (Charon *et al*., 2012; Nakamura, 2020). Each periplasmic flagellum is attached subterminally to one end of the cell poles and extends toward the other end. Spirochete species vary significantly in the number of periplasmic flagella and whether the flagella overlap in the center of the cell (Izard et al., 2009; Liu et al., 2010; Raddi et al., 2012; Wunder et al., 2016; Zhang et al., 2020).

Like the external flagella of the model organisms *Escherichia coli* and *Salmonella enterica*, the periplasmic flagellum in spirochetes consists of a motor, hook, and filament (Chang et al., 2019; Charon *et al*., 2012). The motor is a rotary machine responsible for the assembly and function of the periplasmic flagellum. Most components of the spirochetal flagellar motor have highly conserved counterparts in the external flagellar motor: the MS ring, C ring, rod, export apparatus, and stator (Carroll and Liu, 2020; Chen et al., 2011; Zhao et al., 2014). Uniquely, a spirochete-specific flagellar component – termed collar – not only contributes to the distinct spirochetal motor structures but also plays a role in recruiting 16 torque-generating stator units (Izard *et al*., 2009; Kudryashev et al., 2009; Liu *et al*., 2010; Liu et al., 2009; Moon et al., 2016; Moon et al., 2018; Murphy et al., 2006; Raddi *et al*., 2012; Xu et al., 2020), presumably enabling the increased torque required for spirochetes to swim through complex viscous host environments (Beeby et al., 2016). The collar structure also contributes to make the spirochetal flagellar motor considerably larger and more complicated than its counterparts in *E. coli* and *S. enterica* (Zhao *et al*., 2014). However, how the collar supports the production of high torque by the spirochetal flagellar motor has remained poorly understood.

*B. burgdorferi* has emerged as an ideal model system for understanding the unique structure and function of periplasmic flagella (Chang and Liu, 2019; Charon *et al*., 2012). At each cell pole, 7-11 periplasmic flagella wrap inward as a flat ribbon along the cell body and overlap in the middle of the cell (Fig. 1A)(Charon et al., 2009; Zhang *et al*., 2020). A combination of genetic and cryo-electron tomography (cryo-ET) approaches has enabled *in-situ* visualization of *B. burgdorferi* flagellar motors at an unprecedented resolution, unveiling unique features of this complex machine (Carroll and Liu, 2020; Chang and Liu, 2019). Specifically, comparative analyses of wild type, stator deletion mutant Δ*motB*, and collar deletion mutant Δ*flbB* provided direct evidence that the collar is important for stator assembly, flagellar orientation, cell morphology, and motility in *B. burgdorferi* (Chang *et al*., 2019; Moon *et al*., 2016; Moon *et al*., 2018). The collar is a large complex consisting of the inner core and the outer, turbine-like structure. Three collar proteins have been identified in *B. burgdorferi* (Moon *et al*., 2016; Moon *et al*., 2018; Xu *et al*., 2020): FlbB (BB0286) appears to serve as the base of the collar structure (Moon *et al*., 2016), BB0236 is involved in collar assembly (Moon *et al*., 2018), and FlcA (BB0326) forms the turbine-like structure, directly interacting with the stator units (Xu *et al*., 2020). Given that the overall structure of the collar is 79 nm in diameter and ∼20 nm in height, additional proteins are likely involved in collar assembly. Moreover, the large collar structure must be flexible to accommodate the highly curved membrane at the cell tip (Chang *et al*., 2019). How the collar assembles and contributes to stator assembly is essential for understanding the unique spirochete motility.

**Figure 1.**
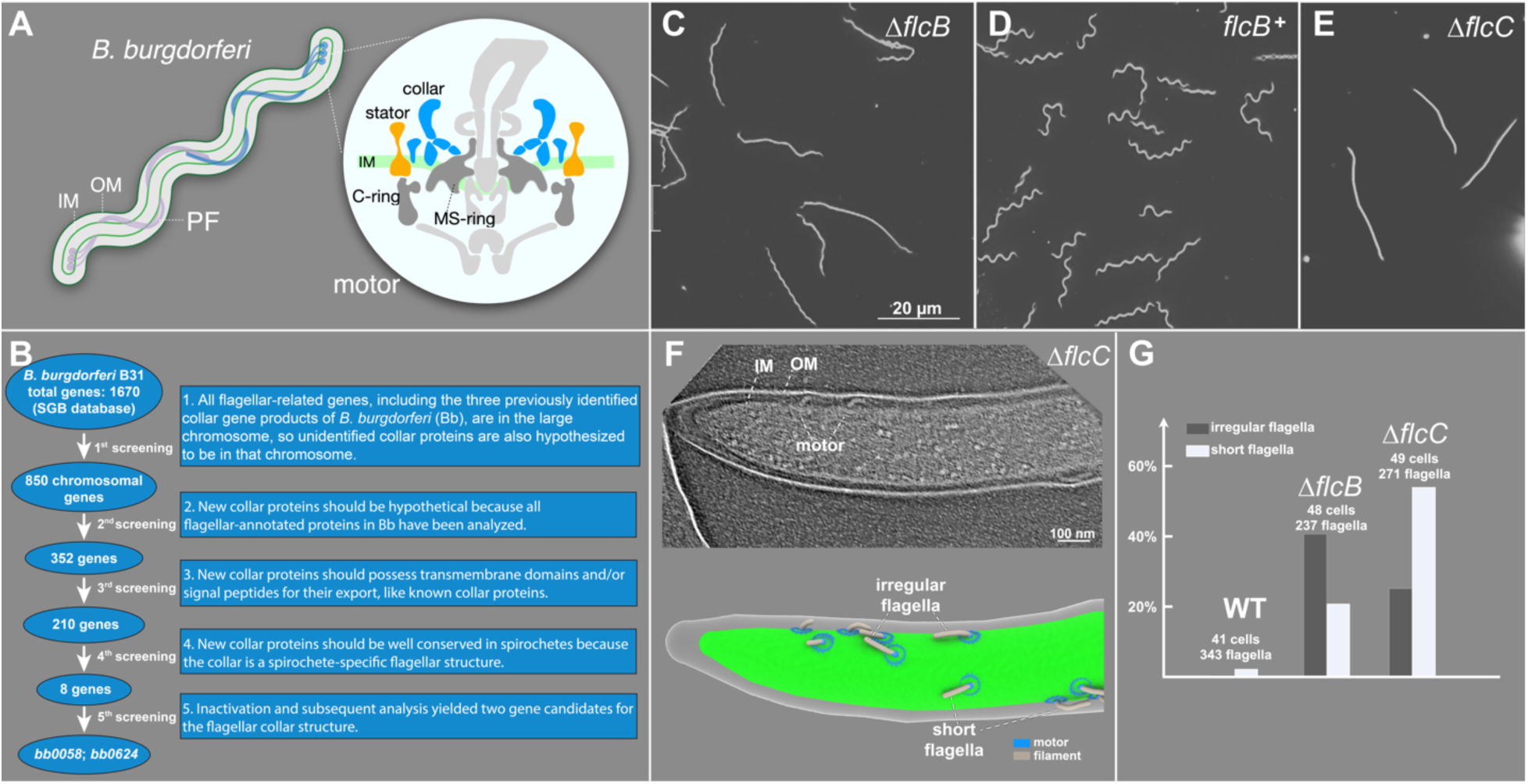
BB0058 (FlcB) and BB0624 (FlcC) are potential collar proteins in *B. burgdorferi*. (A) Schematic models of the periplasmic flagellum (PF) and motor in a *B. burgdorferi* cell. (B) Based on the common features of the previously identified flagellar collar proteins, all 1670 gene-encoded proteins of *B. burgdorferi* were manually screened from the Spirochetes Genome Browser (http://sgb.leibniz-fli.de/cgi/list.pl?sid=24&c_sid=yes&ssi=free). Transmembrane domains and signal peptides were predicted using TMHMM, Phobius, and SignalP-5.0 programs. Subsequent screenings resulted in identification of eight potential candidate collar proteins. (C-E) Dark-field microscopic images showing the characteristic rod-shaped morphology of Δ*flcB* mutant cells, flat-wave morphology of complemented *flcB*^*+*^ cells, and rod-shaped morphology of Δ*flcC* mutant cells, respectively. (F) A representative tomographic slice of the Δ*flcC* cell tip (top) and corresponding 3D surface view (bottom), showing the irregular and short flagella in the mutant cell. (G) Statistical analysis of the flagellar phenotype in WT, Δ*flcB*, and Δ*flcC* cells. A normal flagellum is defined as being oriented toward the other pole of the cell body. An abnormal or irregular periplasmic flagellum is defined as being tilted toward the cell pole from where it originated. The total number of cells and periplasmic flagella analyzed for each strain is shown at the top of the corresponding column.

In this study, we identify two novel collar proteins, FlcB (BB0058) and FlcC (BB0624), each responsible for distinct portions of the collar. Together with studies of other collar proteins (Moon *et al*., 2016; Moon *et al*., 2018; Xu *et al*., 2020), our high-resolution *in-situ* structural analyses of the *B. burgdorferi* flagellar motor provide a molecular basis for the assembly and flexibility of the periplasmic collar complex and its critical roles in the assembly of the stator complexes. Our results also highlight how the collar contributes to the distinct motility that allows spirochetes to swim through complex environments, such as inside ticks and vertebrate hosts.

## Results

### BB0058 and BB0624 are potential collar proteins

To better understand collar assembly and function, we devised a subtractive bioinformatic approach to identify eight potential collar proteins (Fig. 1B). Each corresponding mutant was constructed and analyzed with respect to motility and morphology phenotypes (Fig. S1 and S2). Two of these (*bb0058* and *bb0624*) were ultimately identified as the genes encoding potential collar proteins for the following reasons: 1) Δ*bb0058* and Δ*bb0624* mutant cells exhibited rod-shaped morphology instead of the characteristic flat-wave morphology in wild-type (WT) spirochetes (Fig. 1C, E). Δ*bb0058* mutant cells were significantly less motile than WT cells, whereas the Δ*bb0624* mutant cells were completely non-motile (Fig. S3). These mutants exhibited no polar effects on downstream gene expression. A complemented Δ*bb0058* mutant *in cis* (*bb0058*^*+*^) was constructed as described (Pitzer *et al*., 2011), and it restored the morphology and motility phenotypes to WT levels (Fig. 1C, D and S3). 2) Domain analysis data suggest that BB0058 possesses multiple tetratricopeptide repeat (TPR) domains, and both BB0058 and BB0624 possess a signal peptide at their N-terminal region that is likely required for their export across the membrane (not shown in current work). 3) Cryo-ET reconstructions of the cell tips indicate that the Δ*bb0058* and Δ*bb0624* cells possess approximately 40% and 34% fewer flagella than WT cells (Fig. 1G), respectively. In addition, the flagella in both Δ*bb0058* and Δ*bb0624* cells appear to show shorter lengths and abnormal orientations (Fig. 1F, G), with filaments extending toward their pole of origin instead of toward the other cell pole, as in WT cells. Similar shorter length and abnormal orientation phenotypes were also observed in our previously reported collar gene mutants (Xu *et al*., 2020).

### FlcB is a novel flagellar protein that contributes to the middle portion of the collar

To determine whether BB0058 is involved in assembly of the collar complex, we used cryo-ET and sub-tomogram averaging to resolve the *in-situ* structures of the flagellar motor in Δ*bb0058* and *bb0058*^*+*^ cells. Compared to the WT motor (Fig. 2A, E), a bridge-like structure near the interface between the collar and the MS ring is absent from the Δ*bb0058* motor (Fig. 2B, F), but this structure is restored in the complemented *bb0058*^*+*^ motor (Fig. 2C, G), suggesting that BB0058 is responsible for the formation of the bridge-like structure of the collar (Fig. 2D, H). We therefore renamed BB0058 as periplasmic <u>fl</u>agellar <u>c</u>ollar protein <u>B</u> (FlcB). Notably, in the spirochetal flagellar motor, sixteen copies of this bridge-like structure form the FlcB ring directly above the MS ring (Fig. 2D, H). The FlcB ring does not directly interact with the stator complexes or the MS ring yet has a significant impact on flagellar rotation and bacterial motility.

**Figure 2.**
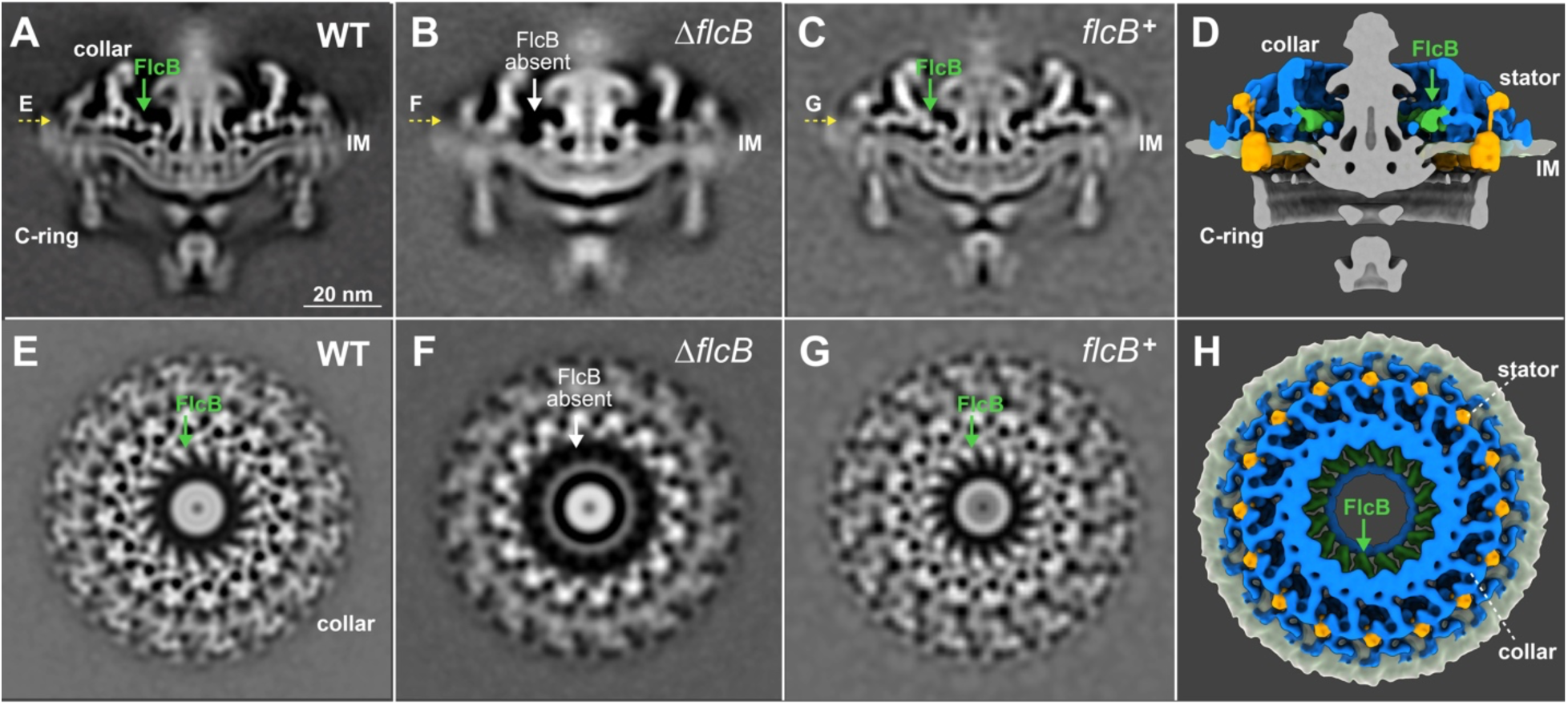
Δ*flcB* (Δ*bb0058*) mutant cells show defects in the flagellar collar structure. (A-C) A central section of the sub-tomogram averages (16-fold symmetrized) of the WT, Δ*flcB*, and *flcB*^+^ flagellar motors, respectively. The middle portion of the collar is absent in the Δ*flcB* motor. (E-G) A top view corresponding to the motor structures shown in A-C (indicated by yellow arrows), respectively. (D, H) A cross and top view of the 3D rendering of the WT flagellar motor, respectively. The FlcB protein is shown in green. Only the collar, stator, and IM are shown in H.

### FlcC is a novel flagellar protein responsible for collar and stator assembly

To identify specific roles of the BB0624 protein, we determined the *in-situ* structure of the Δ*bb0624* motor by cryo-ET and sub-tomogram averaging, revealing that the top portion of the collar is absent (Fig. 3B). BB0624 is therefore a collar protein, renamed hereafter as periplasmic <u>fl</u>agellar <u>c</u>ollar protein <u>C</u> (FlcC). Furthermore, the densities corresponding to the stator complexes in the Δ*flcC* motor are considerably different from those in the WT motor, suggesting that FlcC directly impacts not only collar formation but also stator assembly. To estimate stator complex numbers in WT and these two new mutants, focused alignment and classification were utilized to analyze the stator densities. For the Δ*flcC* mutant, the class with stator density (Fig. 3C) accounts for ∼40% of the total collar subunits, while the class without stator density (Fig. 3D) accounts for ∼60%, indicating that stator occupancy in the Δ*flcC* motor is ∼40%, considerably lower than in the WT (∼96%) and Δ*flcB* motors (∼94%). This result is consistent with immunoblotting data showing that the stator protein MotB is significantly reduced in the Δ*flcC* mutant compared to WT cells (Fig. S2C). Collectively, these results support the model that FlcC functions as a major collar protein directly involved in collar formation and stator assembly.

**Figure 3.**
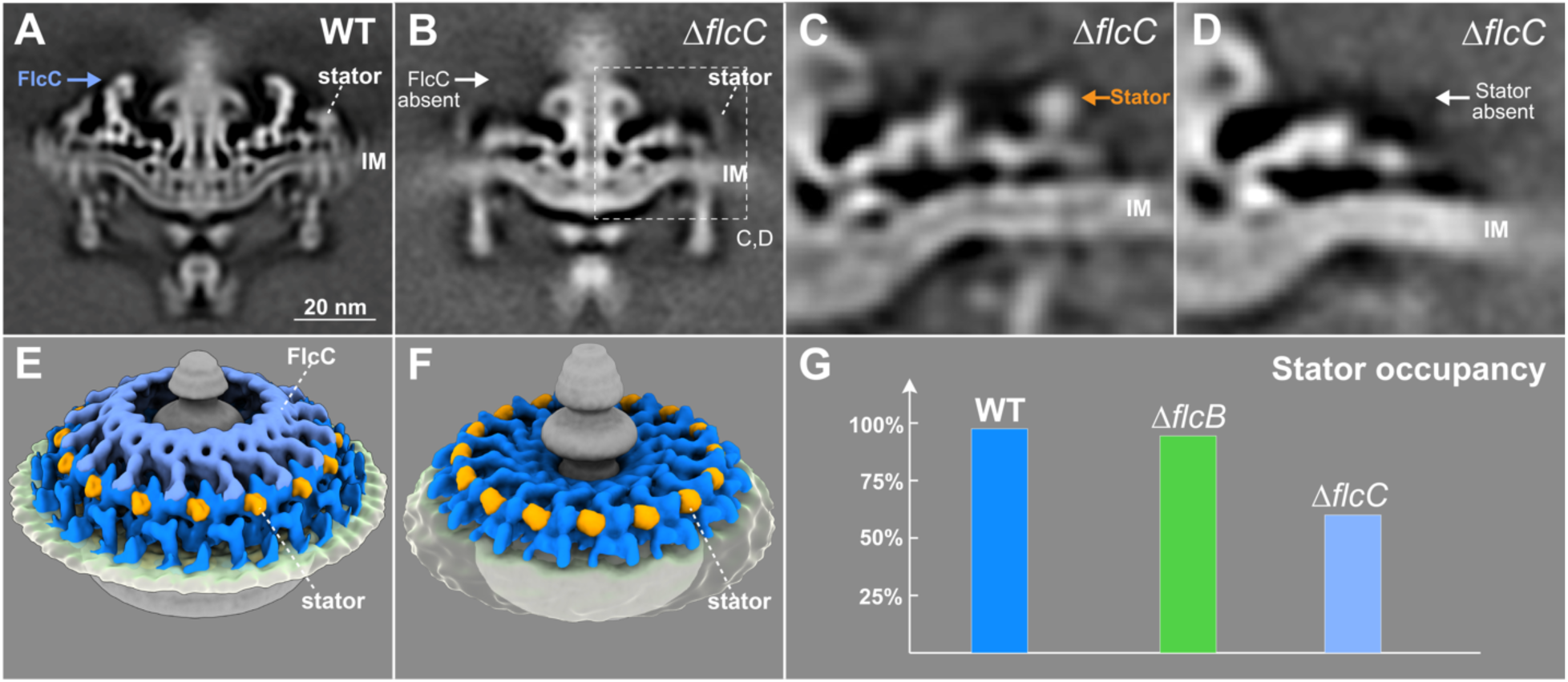
Δ*flcC* mutant cells show defects in the collar structure and have fewer stator units assembled in the motor. (A, B) A central section of the sub-tomogram average (16-fold symmetrized) of WT and Δ*flcC* flagellar motors, respectively. The top portion of the collar (indicated by white arrow) is absent from the Δ*flcC* motor. (C, D) Class averages of the collar region (dash frame indicated in B) with and without the density of the stator complex in the Δ*flcC* motor, respectively. (E, F) A tilted side view of the 3D rendering of WT and Δ*flcC* flagellar motors (with stator complexes), respectively. (G) A histogram showing stator occupancy in the WT, Δ*flcB*, and Δ*flcC* motors, respectively. Refer to the Materials and Methods section for details about the calculation of stator occupancy.

### The molecular architecture of the collar reveals its intrinsic plasticity

Five collar proteins have been identified and characterized in *B. burgdorferi*: FlbB (Moon *et al*., 2016), BB0236 (Moon *et al*., 2018), FlcA (Xu *et al*., 2020), FlcB (BB0058), and FlcC (BB0624). To understand how these proteins assemble as the complex collar, we developed a sophisticated approach to analyze the collar structure in the absence of the stator complexes. First, we generated an asymmetric reconstruction of the Δ*motB* motors (Fig. S4A, B) (Chang *et al*., 2019). The asymmetric reconstruction reveals sixteen collar subunits and their associated membrane curvature (Fig. S4C), consistent with the observation that the motors are embedded in a highly curved membrane cylinder. Second, to determine the collar subunit structure at higher resolution, we extracted 16 collar subunits from each motor and performed 3D classification and focused refinement (Fig. S4D, E, Movie S1). Third, the high-resolution structure of the collar subunit was then mapped back to the asymmetric reconstruction of the Δ*motB* motor structure to obtain a detailed overview of the collar complex (Fig. 4). The exact location of each collar protein was defined by comparing the high-resolution *in-situ* structure of the Δ*motB* motor with specific collar mutant structures reported in this study (Fig. 2, 3) and previously (Moon *et al*., 2016; Moon *et al*., 2018; Xu *et al*., 2020) and analyzing the protein-protein interaction data (Fig. S5). In the large collar complex (79 nm in diameter) (Fig. 4A, B), FlcA is closely associated with the membrane and forms the turbine-like structure (Fig. 4). FlcB forms a ring at the base of the collar (Fig. 4A, B). FlcC is located on top of the collar structure (Fig. 4A, C, D), and FlbB is located approximately in the middle of the collar complex (Fig. 4B). Although some components of the collar remain undefined, it is evident that the collar complex is composed of multiple different proteins, each contributing to a distinct portion of the highly modular, flexible architecture of the collar complex. Importantly, this highly modular architecture of the collar enables extensive remodeling to accommodate the curvature of the membrane cylinder, which is ubiquitous in spirochetes and other bacteria (Fig. 4C, D).

**Figure 4.**
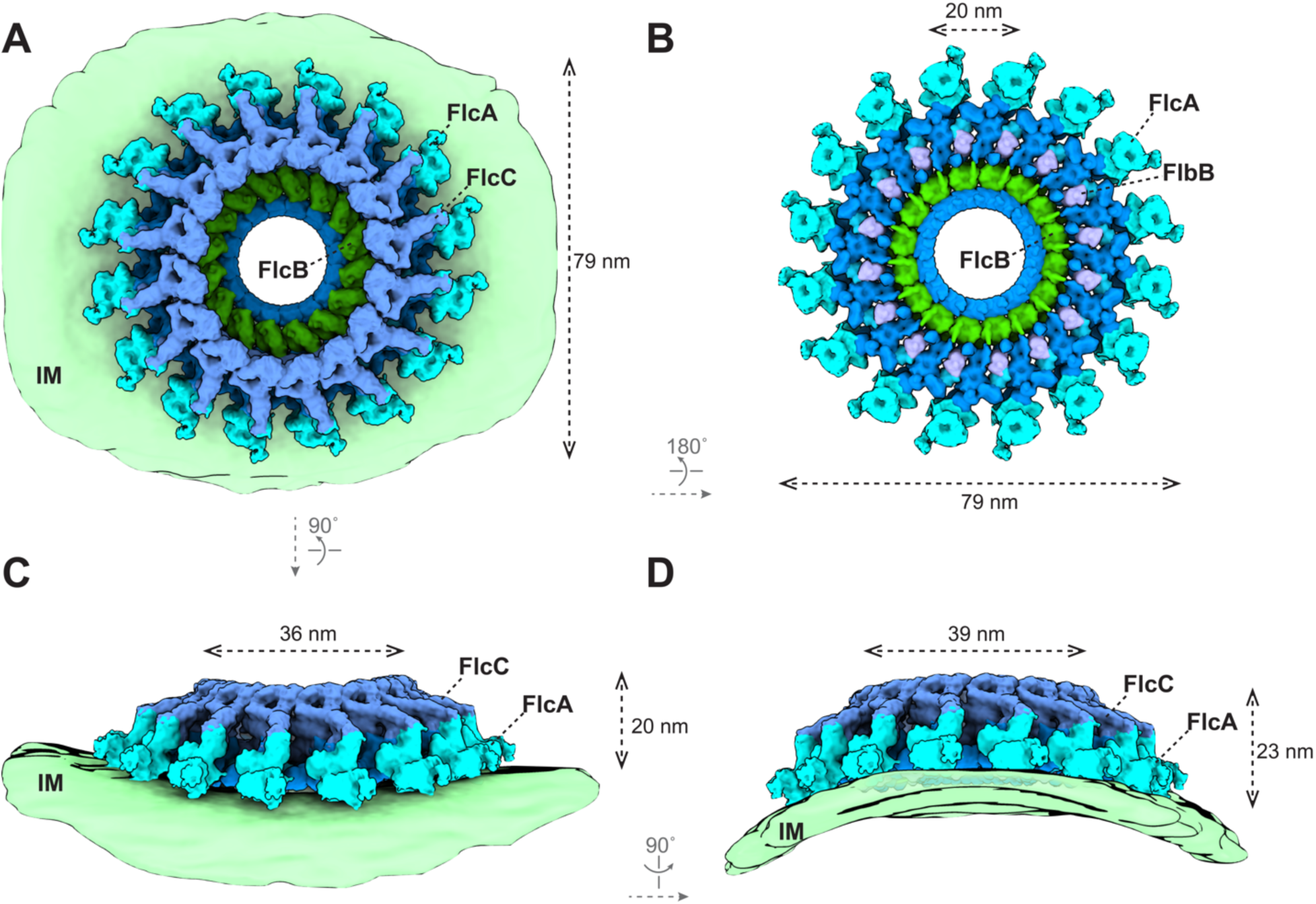
Structure of the flagellar collar in Δ*motB B. burgdorferi*. (A-D) 3D surface view of the whole collar complex in Δ*motB B. burgdorferi*. The collar is ∼79 nm in diameter and embeds on the IM. Each known collar protein has 16 copies and assembles into a ring structure. The membrane of the spirochetal cells shows a clear curvature and distorts the collar complex (C, D). The height of the collar is 20 nm on the flat IM (C) and 23 nm on the curved IM (D). The inner diameter of the FlcC ring is 36 nm on the flat IM (C) and 39 nm on the curved IM (D). The FlcA proteins are colored in ice, FlcB in green, FlcC in light blue, and FlbB in purple. The unknown collar proteins are colored in dark blue.

### The collar facilitates the assembly and function of the stator complexes

The collar is important for stator assembly and recruitment in spirochetes (Moon *et al*., 2016). To determine the detailed interactions between the collar and stator complexes, we analyzed WT motors (Fig. S4G-K) and determined the high-resolution asymmetric reconstruction of the collar (Fig. 5). The overall size and shape of the collar remains similar in both WT and Δ*motB* motors (Fig. 4, 5). In the WT motor, 16 stator units are closely associated with the collar complex through multiple interactions. The periplasmic domain of the stator complex directly interacts with FlcA and additional unknown collar proteins (Fig. 5 and Fig. S4J), likely stabilizing its assembly around the motor. Furthermore, 16 stator complexes appear perfectly embedded in the curved membrane cylinder around the motor. By contrast, the stator complexes fail to assemble around the motor in the absence of the entire collar or its periphery (Moon *et al*., 2016; Moon *et al*., 2018; Xu *et al*., 2020). Therefore, the collar has evolved a remarkable modular architecture ideal not only to recruit the stator complexes but also to stabilize the stator ring with 16 complexes around the motor, thus ensuring maximal torque generation (Fig. 5).

**Figure 5.**
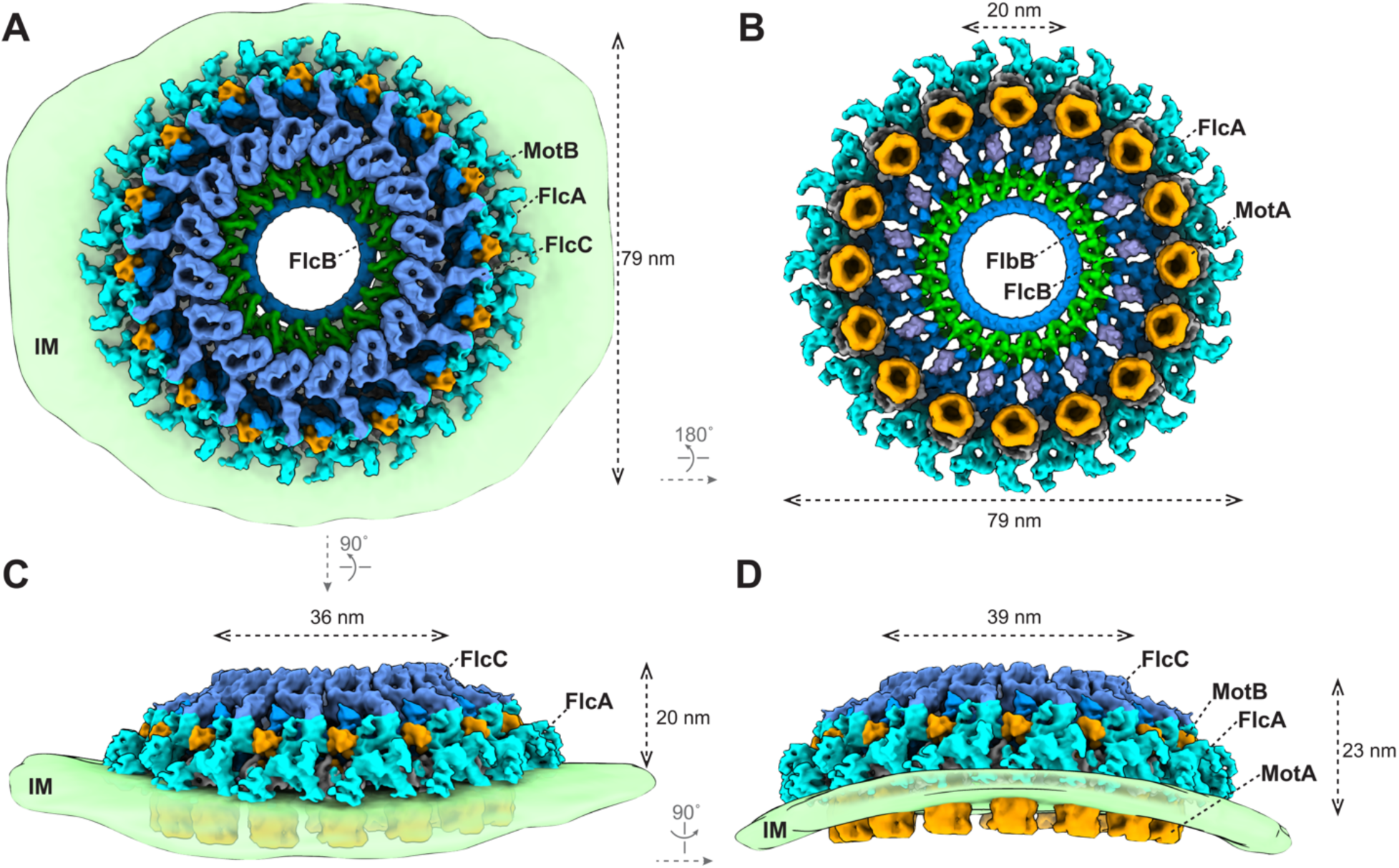
The collar facilitates the assembly of 16 stator complexes in a highly curved membrane. (A-D) 3D surface view of the whole collar complex in WT *B. burgdorferi*. The diameter and height of the collar are indicated. The curvature of the IM is clearly resolved in the asymmetric reconstruction of the motor structure (C, D). 16 stator units embed on the IM and are surrounded by the collar proteins. FlcA directly interacts with the stator units. The color scheme is the same as in Fig. 4.

## Discussion

Spirochete motility is unique among bacteria, due to the location and distinct assembly of periplasmic flagella. It is increasingly evident that the periplasmic flagellum possesses a unique multi-protein collar important for the assembly of periplasmic flagella and motility of spirochetes. Five spirochete-specific collar proteins in *B. burgdorferi* are involved in collar assembly. Given that these collar proteins are well conserved in spirochetes, their homologs are likely involved in collar assembly across diverse spirochetes (Chen *et al*., 2011; Zhao *et al*., 2014).

One of the most remarkable features of *B. burgdorferi* is that multiple flagellar motors are embedded in the inner membrane cylinder in a highly organized pattern (Fig. 6A). As a result, the cell cylinder diameter varies remarkably, ranging from ∼100 nm to ∼300 nm (Fig. 6A). The collar must therefore be highly flexible to accommodate variable membrane curvatures (Fig. 6B, C, Movie S2). Indeed, our studies have clearly demonstrated that the collar has a highly modular architecture due to the highly coordinated assembly of multiple spirochete-specific proteins (including several transmembrane proteins). This modular architecture may be of key importance for facilitating the remarkable plasticity of the collar. Moreover, multiple collar proteins directly interact with the stator complexes. Therefore, the unique plasticity of the collar also facilitates the recruitment and stabilization of maximal numbers of stator complexes around the motor even in highly curved membrane environments (Fig. 6B, C, Movie S2). That the entire flagellar motor remodels and adapts to accommodate variable membrane environments is crucial to generate the highest torques required to constantly drive the motility of spirochetes and benefit their distinct lifestyle.

**Figure 6.**
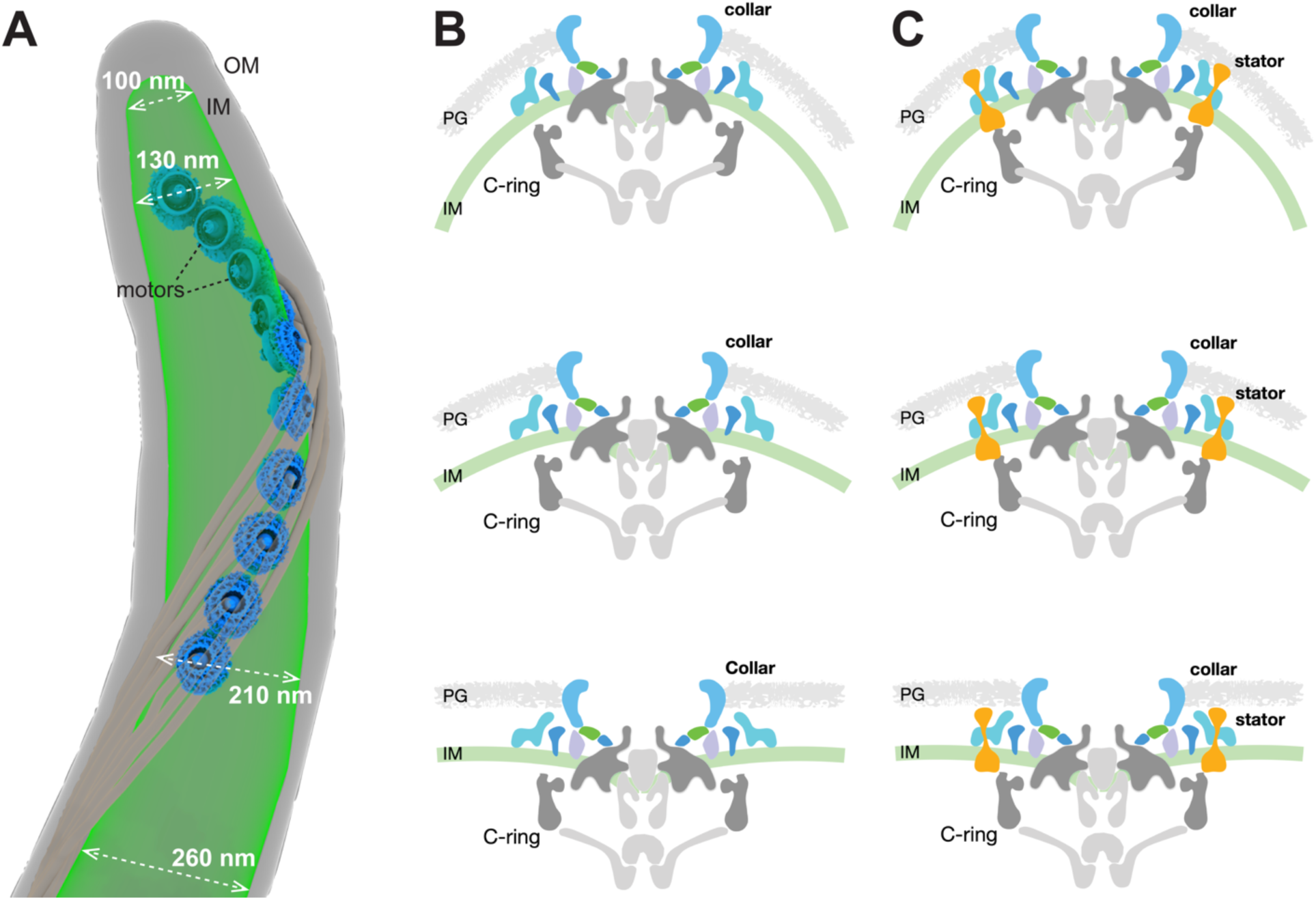
The mosaic collar complex changes conformation to accommodate the inner membrane curvature in *B. burgdorferi*. (A) Surface rendering of a representative WT cell tip. Ten motors embed in the IM and distribute along the cell tip. The diameter of the IM at different positions at the cell tip is indicated. Note that the IM is more curved (has a smaller diameter) at the positions closer to the cell tip. (B, C) The flagellar motor embeds on the IM and is distorted due to the IM curvature. The periplasmic collar complex surrounds the stator units and can change conformation to fit the IM curvature, facilitating the assembly and function of spirochetal motors in highly curved membrane environments.

In summary, we have identified and characterized multiple novel collar proteins in *B. burgdorferi*, providing a molecular basis for understanding the remarkable structural plasticity of this multi-protein spirochetal complex. The collar not only enables the assembly of the motor in the curved membrane of spirochetes but also provides a structural scaffold for stator recruitment and stabilization, both essential for the function of periplasmic flagella and motility in spirochetes. Identification of additional collar proteins based on the protein-protein interactions, along with high-resolution *in-situ* structural analyses, will provide further insights into how the structural plasticity of the collar is required for motility in spirochetes.

## Materials and methods

### Bacterial strains and growth conditions

High-passage *B. burgdorferi* strain B31-A was used as the wild-type (WT) clone throughout this study (Bono *et al*., 2000; Elias et al., 2002). The mutant Δ*bb0058* (Δ*flcB*), Δ*bb0624* (*ΔflcC*), and complemented *flcB*^+^ strains were constructed as described below. *B. burgdorferi* cells were cultivated in liquid Barbour-Stoenner-Kelly (BSK-II) broth or agarose plates and incubated at 35°C in a 2.5% CO_2_ incubator, as reported previously (Motaleb *et al*., 2007; Sultan *et al*., 2013). Antibiotics, when required, were included in the *B. burgdorferi* medium at the following concentrations: 200 μg/ml kanamycin and/or 100 μg/ml streptomycin. *Escherichia coli* strains were grown at room temperature or 37°C in liquid Luria-Bertani (LB) broth or plated on LB agar (Bertani, 1951; 2004). When required, 100 μg/ml ampicillin, 100 μg/ml spectinomycin, 0.2% glucose, 80 μg/ml 5-bromo-4-chloro-3-indolyl-β-D-galactopyranoside (X-gal), and/or 0.5 mM isopropyl β-d-1-thiogalactopyranoside (IPTG) was supplemented into LB medium.

### Bioinformatics

Basic local alignment search tool (BLAST) (Altschul *et al*., 1990; Altschul *et al*., 1997) was utilized to determine protein or gene homologs from the sequence database. The lower the E-value (lower than 0), the more significant the score is. Signal peptides were predicted using SignalP 5.0 and Phobius programs (Almagro Armenteros *et al*., 2019; Kall *et al*., 2005; 2007). Protein domains were analyzed using the Conserved Domain Database (Marchler-Bauer *et al*., 2017; Marchler-Bauer *et al*., 2015) and Pfam (El-Gebali *et al*., 2019; Sonnhammer *et al*., 1997).

### Overexpression of recombinant proteins in *E. coli*

To express the *B. burgdorferi* BB0058 (FlcB) protein in *E. coli*, a DNA fragment harboring the BB0058 open reading frame (ORF) without the signal peptide region (1-20 aa) was PCR-amplified from chromosomal DNA of *B. burgdorferi* B31-A cells using primers PF MBP0058_BamHI (CGTCGAC<u>GGATCC</u>GATACTACAGCATTAGGACATTATC) and PR MBP0058_PstI (TTAATTAC<u>CTGCAG</u>TTATCTTTTTATAAGCACAGTGGCTC) (restriction sites are underlined) and cloned into the pMAL c5x (NEB Inc.) using *Bam*HI and *Pst*I restriction sites to produce the MBP-BB0058 protein. MBP-MotB was similarly generated. In brief, the coding sequence of MBP from pMAl c5x was fused to the 3’ end of the coding region of MotB without its transmembrane domain, aa 1-44 using PCR, and then cloned into pET28a(+) (Novagen Inc.). Similarly, 1xFLAG (DYKDDDDK) tagged BB0624 (FlcC) and FlcA-C (BB0326-C) were constructed for affinity blotting. In brief, 1xFLAG tag coding sequence (GACTACAAAGACGATGACGACAAG) was fused to the coding regions of BB0624 without the signal peptide region, aa 1-22 and the C-terminus aa 360-931 of BB0326 using PCR amplification with primers PF MBP0624_BamHI (CGTCGAC<u>GGATCC</u>GATTACAAGGGCCTTGATTTTAAAATC) and PR MBP0624FLAGc_PstI (TAATTAC<u>CTGCAG</u>TTACTTGTCGTCATCGTCTTTGTAGTCCTTTTCCTTAATGCCAGT ATTTTG), PF HisThroBB326C572aa_NdeI (GGCAGC<u>CATATG</u>GCCTCTGAGAGCAAGTATAAAGAG) and PR PR BB0326C572aaFLAGc_NotI (GACGATATC<u>GCGGCCGC</u>TTACTTGTCGTCATCGTCTTTGTAGTCAAGTTTTTCGGAT AAATTTTC), respectively, which were then cloned into pMAL c5x (NEB Inc.). Expression of MBP tag, MBP tagged MCP5, FliL, FlbB, BB0236, and FlcA-C and 1xFLAG tagged MotB, FlbB, BB0236, and FliL were described elsewhere (Moon *et al*., 2016; Moon *et al*., 2018; Xu *et al*., 2020).

All *E. coli* strains were induced with 0.5 mM IPTG at room temperature, and purifications of recombinant proteins were performed using amylose resin for MBP-tagged proteins and HisPure Ni-NTA resin for His-tagged proteins.

### SDS-PAGE, immunoblot, and affinity blotting

Sodium dodecyl sulfate polyacrylamide gel electrophoresis (SDS-PAGE) was performed as described (Li *et al*., 2002; Motaleb *et al*., 2000; Sultan *et al*., 2013). Exponentially growing *B. burgdorferi* cells were harvested and washed with phosphate-buffered saline (PBS) and resuspended in the same buffer to process the preparation of cell lysate for SDS-PAGE. Immunoblotting with *B. burgdorferi* FlaB, MotB, FlbB, FliL, FliG1, and DnaK-specific antibodies (Barbour *et al*., 1986; Carroll *et al*., 2001; Coleman and Benach, 1992; Li *et al*., 2010; Moon *et al*., 2016; Motaleb *et al*., 2011a) was performed using Pierce™ enhanced chemiluminescence western blotting substrate (Thermo Fisher Scientific Inc.). Protein concentrations were determined using a Bio-Rad protein assay kit with bovine serum albumin as the standard. Unless specified, approximately 10 µg of cell lysates were subjected to SDS-PAGE.

Far western or affinity blot assays were performed as described previously (Kariu *et al* l., 2015; Moon *et al*., 2016; Moon *et al*., 2018; Toker and Macnab, 1997; Xu *et al*., 2020). In brief, 1 µg purified recombinant proteins was subjected to SDS-PAGE and transferred to polyvinylidene difluoride membranes. The membranes were blocked in the blocking solution (5% skim milk, 10mM Tris, 150mM NaCl and 0.3% Tween 20, pH 7.4) with gentle shaking for 4 to 6 hours at room temperature and then incubated with purified 1xFLAG tagged protein at concentration 2 µg/ml in blocking solution overnight. The membranes were washed 3 times with washing buffer (10mM Tris, 150mM NaCl and 0.3% Tween 20, pH 7.4) and then probed with monoclonal anti-FLAG^®^ M2 antibody (Sigma-Aldrich Co. LLC).

### Construction of the *bb0058* and *bb0624* mutants and *bb0058* complemented strain

Construction of the *bb0058* (*flcB*) and *bb0624* (*flcC*) inactivation plasmid, electroporation, and plating of *B. burgdorferi* were performed as described earlier (Motaleb *et al*., 2000; Novak *et al*., 2016; Sultan *et al*., 2013; Sultan et al., 2010). *bb0058* and *bb0624* were inactivated individually using a promoterless kanamycin resistance cassette (*Pl-Kan*), as reported in detail (Sultan *et al*., 2010). The Δ*bb0058* mutant strain was complemented *in cis* by chromosomal integration using the pXLF14301 suicide vector, as described (Li *et al*., 2007; Pitzer *et al*., 2011). In brief, the native promoter regions of the *bb0058* (*P*_*bb0061*_) and *bb0058* genes were separately PCR-amplified from WT *B. burgdorferi* strain B31-A genomic DNA using primer pairs PF Pbb0061bb0058SpeI (TGTCTAGA<u>ACTAGT</u>CCGGCTATTAAATGTTTTTCGCAATC) and P1R Pbb0061bb0058 (AAAAACCAATTAAAATTCATATATTTTACATGCCCCCCTA), and P2F Pbb0061bb0058 (TAGGGGGGCATGTAAAATATATGAATTTTAATTGGTTTTT) and PR Pbb0061bb0058NotI (CTCGGGTA<u>GCGGCCGC</u>CTATCTTTTTATAAGCACAGTGGC), and fused together by overlapping PCR to generate *P*_*bb0061*_-*bb0058*, which was inserted into derivative pXLF14301::P_*flgB*_*-Strep* (Promnares *et al*., 2009; Yang *et al*., 2009; Yang *et al*., 2010; Zhang *et al*., 2009) using *SpeI* and *NotI* restriction sites, yielding pXLFbb0058. The plasmid was then electroporated into the Δ*bb0058* mutant cells, followed by selection with both kanamycin and streptomycin. The resistant clones were analyzed by PCR to verify the integration of *P*_*bb0061*_-*bb0058*-*P*_*flgB*_*-Strep* within the intergenic region of *bb0445* and *bb0446*. Multiple attempts to complement the Δ*bb0624* mutant *in cis* or *in trans* were unsuccessful, as it is well known that genetic manipulations in *B. burgdorferi* are challenging (Dresser *et al*., 2009; Hyde *et al*., 2009; Kung *et al*., 2016; Miller *et al*., 2013; Moon *et al*., 2016; Pappas *et al*., 2011; Ramsey *et al*., 2017; Stewart *et al*., 2008; Xu *et al*., 2017).

### Dark-field microscopy and swarm plate assays

Exponentially growing *B. burgdorferi* cells were observed using a Zeiss Axio Imager M1 dark-field microscope connected to an AxioCam digital camera to determine bacterial morphology, as described previously (Motaleb *et al*., 2007; Motaleb *et al*., 2011b). Swarm plate motility assay was also performed using our established protocol (Motaleb *et al*., 2007).

### Cryo-ET data collection and tomogram reconstruction

Frozen-hydrated specimens were prepared as described previously (Liu *et al*., 2009). In brief, various clones of exponentially growing *B. burgdorferi* cells were centrifuged individually at 5,000 x g for ∼5 mins, and the resulting pellets were suspended in PBS to achieve cell concentration ∼1 × 10^8^/ml. After adding 10 nm gold marker solution, 5 µl of the cell suspension was placed on freshly glow-discharged (for ∼25 s) holey carbon grids (Quantifoil Cu R2/1, 200 mesh). The grids were front blotted with Whatman filter paper and rapidly frozen in liquid ethane, using a homemade plunger apparatus as described previously (Liu *et al*., 2009). The grids were then imaged using a 300-kV electron microscope (Titan Krios, Thermo Fisher Scientific) equipped with a field emission gun, a Volta Phase Plate (VPP), and a post-GIF Direct Electron Detector (Gatan K2 Summit or K3 Summit). SerialEM was used to collect all tilt series (Mastronarde, 2005). The defocus was set as close to 0 µm as possible for those tilt series collected with VPP, while the defocus was set ∼-3µm for those collected without VPP. A total dose of ∼80 e^−^/Å^2^ is distributed among 35 (or 33) tilt images covering angles from −51° to 51° (or from −48° to 48°) with a tilt step of 3°.

All recorded images were first motion-corrected using MotionCorr2 (Zheng *et al*., 2017) and then stacked and aligned by IMOD (Kremer *et al*., 1996). For the data collected with VPP, the aligned tilt series were directly used to reconstruct tomograms by weighted back-projection using IMOD or by SIRT reconstruction using Tomo3D (Agulleiro and Fernandez, 2015). For the data collected without VPP, Gctf (Zhang, 2016) was used to determine the defocus of each tilt image in the aligned stacks, and the ‘ctfphaseflip’ function in IMOD was used to do the contrast transfer function (CTF) correction for the tilt images. The tomograms were then reconstructed using IMOD or Tomo3D. The number of tomograms used in this work for each strain is shown in Table S1.

### Sub-tomogram analysis

Bacterial flagellar motors were manually picked from the 6× binned tomograms. The sub-tomograms of flagellar motors were analyzed by i3 software package (Winkler, 2007; Winkler *et al*., 2009). Afterwards, the sub-tomograms were extracted from unbinned tomograms with the refined positions and further binned by 2 or 4 based on the requirement for alignment and classification.

Focused refinement of collar region: each flagellar motor has 16 collar subunits. After alignment for the whole motor structure, the regions around 16 collar subunits were first extracted from each motor, and then we refined the 3D alignment and applied 3D classification based on the density of the collar subunit to remove particles with bad contrast or large distortions to obtain the refined structures. The number of flagellar motor and collar subunits used for sub-tomogram averaging is shown in Table S1.

Measurement of stator occupancy: for the WT, Δ*flcB*, and Δ*flcC* motors, we first performed focused refinement to the collar region, as described previously. Then 3D classification was applied to all collar subunits based on the density of the stator complex. The class averages with density of the stator complex were considered as having assembled stator units, while the class averages without density of the stator complex were considered as having no assembled stator units. The number of collar subunits with stator units was divided by the total number of collar subunits to calculate stator occupancy. Fourier shell correlation (FSC) coefficients were calculated by generating the correlation between two randomly divided halves of the aligned images used to estimate the resolution and to generate the final maps.

### Three-dimensional visualization

UCSF Chimera (Pettersen *et al*., 2004) and ChimeraX (Goddard *et al*., 2018) were used for 3D visualization and surface rendering of sub-tomogram averages of the whole motor or collar subunit. For the 3D surface views of the whole collar complex shown in Fig. 4 and Fig. 5, the surface view of each collar protein in Δ*motB* or WT was first segmented by ChimeraX and then fitted to the collar complex of the asymmetric reconstructed motor structure, using the “fitmap” command in ChimeraX. Segmentations of representative reconstructions from WT and *ΔflcC* cell tips were manually constructed using IMOD (Kremer *et al*., 1996).

## Acknowledgments

We thank Jennifer Aronson for critical reading of the manuscript. We thank Jun He and Shenping Wu for assisting cryo-ET data collection. Y.C. and J.L. were supported by grant R01AI087946; X.H. and M.A.M were supported by R01AI132818 from National Institute of Allergy and Infectious Diseases (NIAID) and National Institutes of Health (NIH). Part of cryo-ET data were collected at Yale CryoEM resource that is funded in part by the NIH grant 1S10OD023603-01A1.

## Supplementary materials

**Table S1.** Data collection parameters and number of tomograms, motors, and collar subunits used for sub-tomogram averaging in the current work.

| strain | Camera | Pixel size (Å) | VPP | tilt range | No. of tomograms | No. of motors | No. of collar subunits used for focus refinement |
| --- | --- | --- | --- | --- | --- | --- | --- |
| WT | K2 | 2.747 | Yes | -51° to 51° | 256 | 1748 | 5232 |
|  |  |  | No |  | 189 | 1202 | 11928 |
| *ΔflcB*<br>(*Δbb0058*) | K2 | 3.483 | Yes | -51° to 51° | 41 | 172 | — — |
|  | K3 | 3.384 |  | -48° to 48° | 132 | 885 | — — |
| *(flcB<sup>+</sup>)*<br>(*bb0058<sup>+</sup>*) | K2 | 1.386 | No | -51° to 51° | 79 | 290 | — — |
| *ΔflcC*<br>(*Δbb0624*) | K2 | 3.483 | Yes | -51° to 51° | 40 | 346 | 5536 |
|  | K3 | 3.384 |  | -48° to 48° | 129 | 798 | 12256 |
| *ΔmotB* | K2 | 2.747 | Yes | -51° to 51° | 233 | 1448 | 9427 |

**Figure S1.**
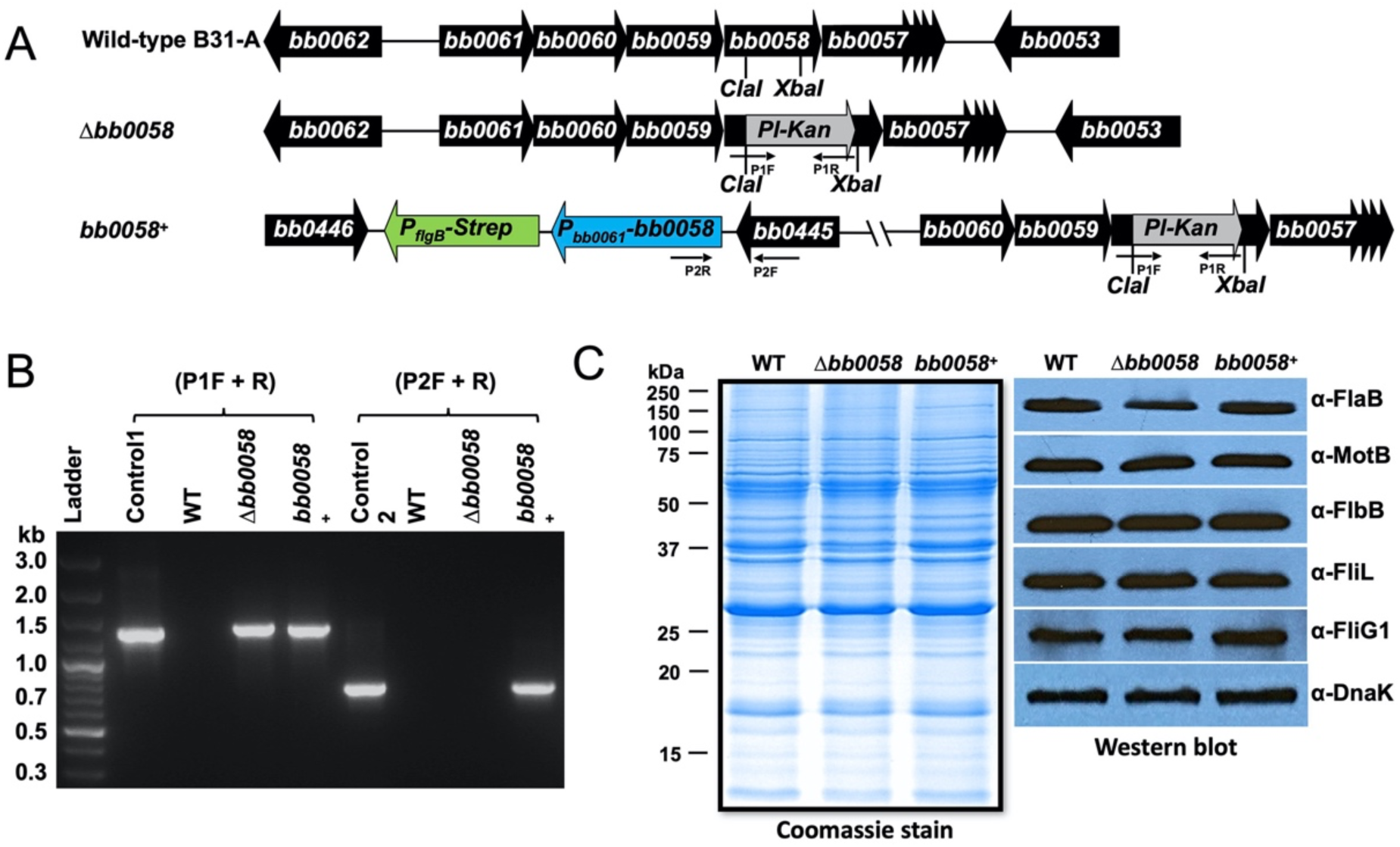
Inactivation, complementation, and determination of flagellar protein synthesis in Δ*bb0058 (*Δ*flcB)*. (A) The *bb0058 (flcB)* gene is located within a putative operon consisting of eight genes (diagram is not to scale). The *bb0058* gene was inactivated by insertion of a promoterless *Pl-Kan* cassette that does not impose any polar effect on downstream gene expression. The mutant was complemented through chromosomal integration. (B) Confirmation of inactivation and complementation of *bb0058* by PCR analysis. (C) Effect of *bb0058* mutation on other flagellar protein synthesis determined by Western blotting. WT, Δ*bb0058*, and *bb0058*^*+*^ cell lysates were subjected to SDS-PAGE followed by Coomassie staining (left) or transferred to a PVDF membrane for immunoblot analysis (right). Immunoblots were performed with *B. burgdorferi* FlaB-, MotB-, FlbB-, FliL-, and FliG1-specific antibodies. DnaK was used as a loading control.

**Figure S2.**
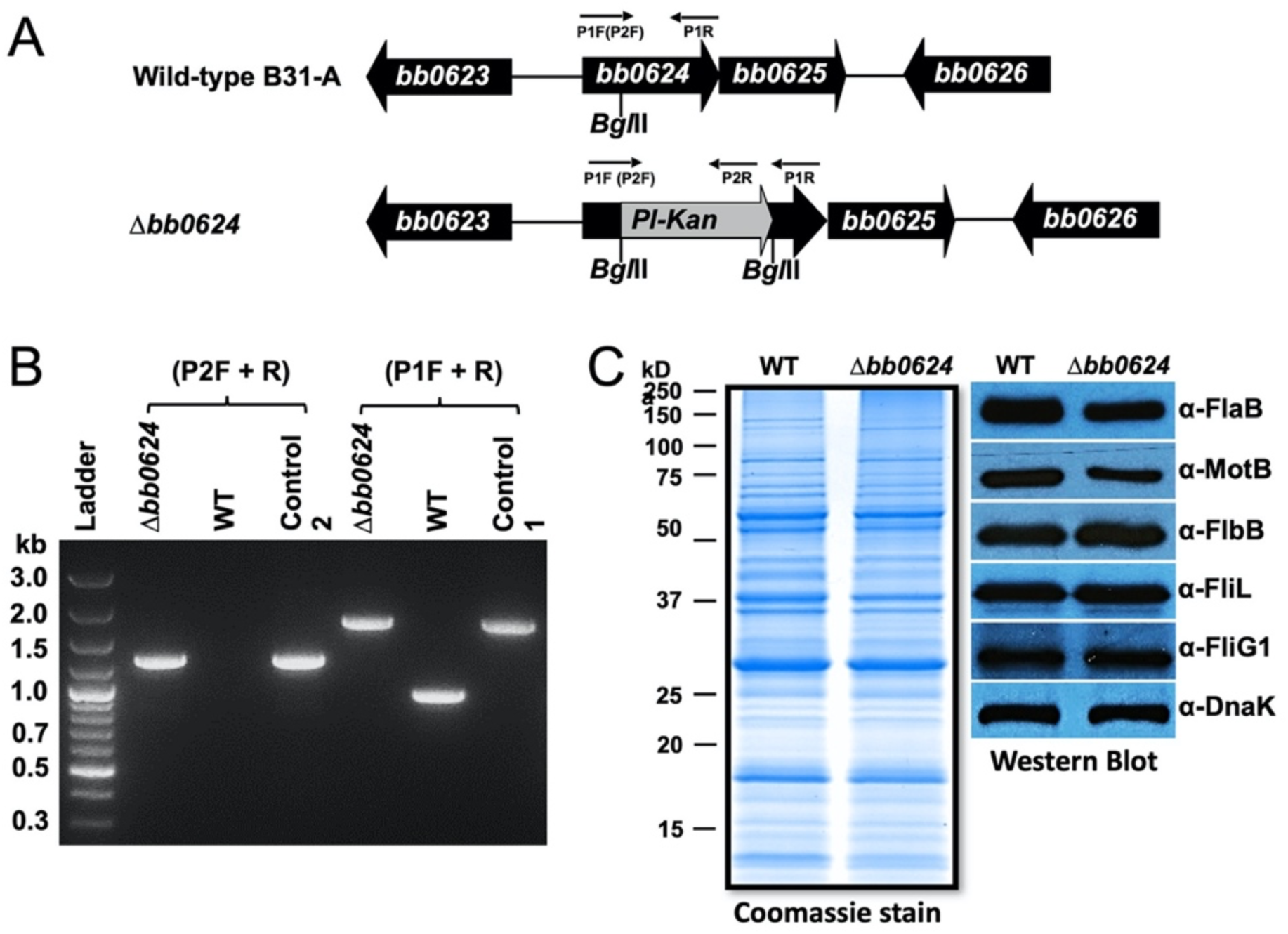
Inactivation and determination of flagellar protein synthesis in Δ*bb0624 (*Δ*flcC)*. (A) The *bb0624 (flcC)* gene is located within a putative operon consisting of two genes (diagram is not to scale). The *bb0624* gene was inactivated by insertion of *Pl-Kan* cassette. (B) Confirmation of inactivation of *bb0624* by PCR analysis. (C) Effect of *bb0624* mutation on other flagellar protein synthesis determined by Western blotting as described above.

**Figure S3.**
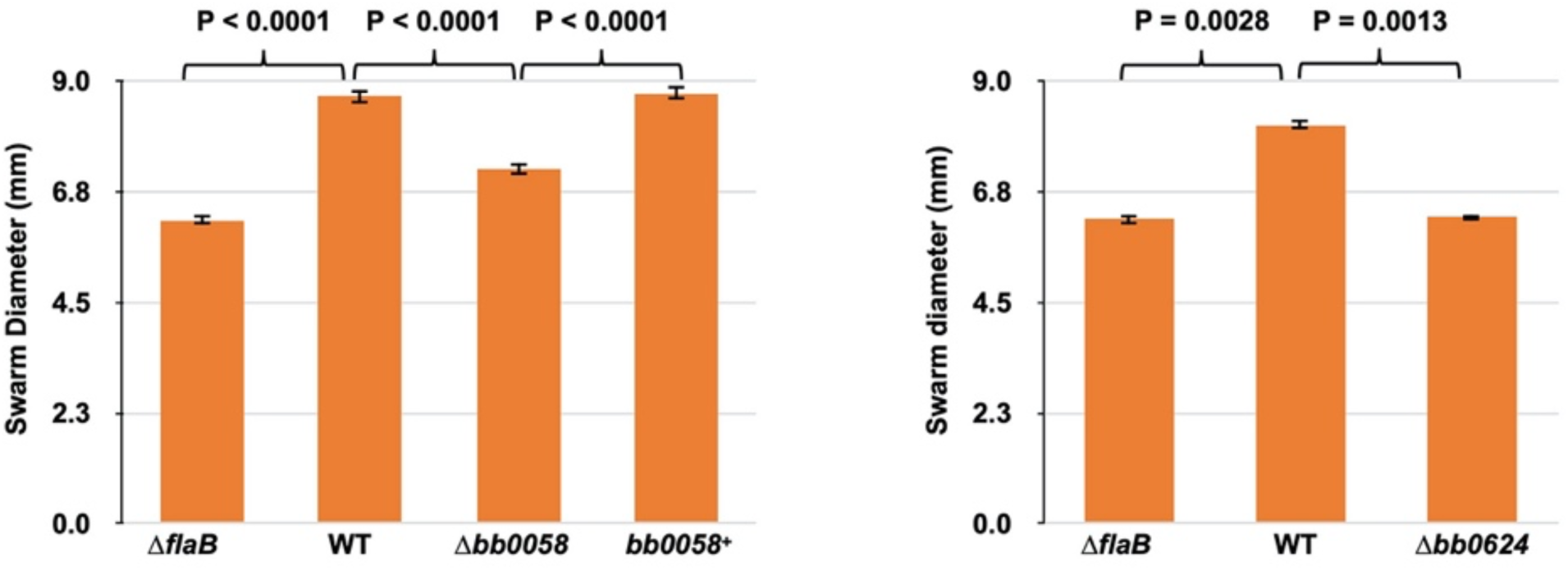
Motility phenotypes of the Δ*bb0058* (Δ*flcB*) and Δ*bb0624* (Δ*flcC*) mutant cells. Swarm plate motility assays of Δ*bb0058* (left) and Δ*bb0624* (right) mutants. Average swarm diameters from three swarm plates are shown at millimeter scale. Non-motile flagellar filament mutant Δ*flaB* was used as control. Bars represent mean ± standard deviation of the mean from three plates. P-values between samples are shown at the top. A P-value of <0.05 is considered different.

**Figure S4.**
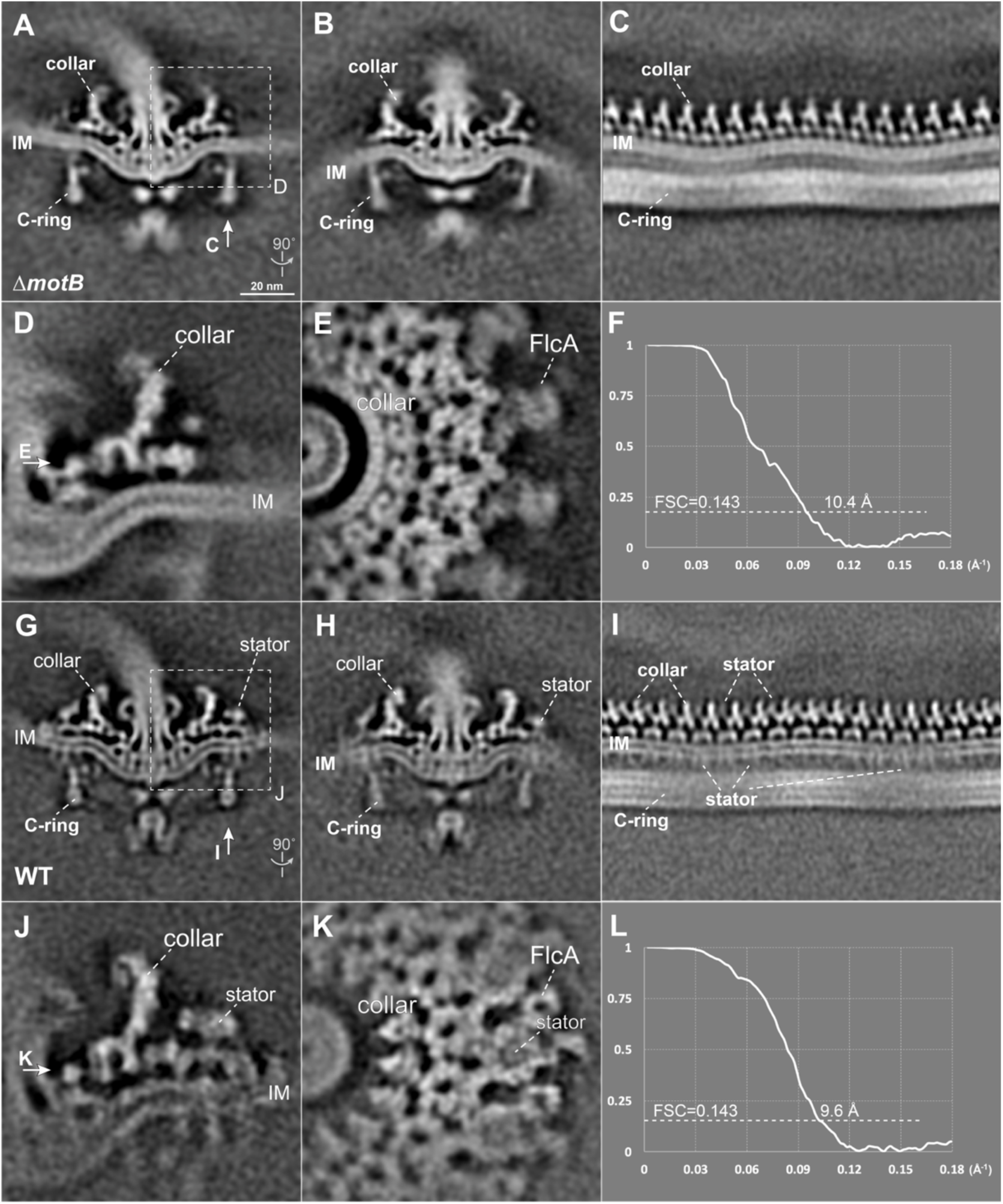
Structures of the flagellar collar in Δ*motB* and WT *B. burgdorferi*. (A, B) Central sections of the asymmetric reconstruction of the Δ*motB* motor. (C) A section from the unrolled map of the Δ*motB* motor showing the collar inserted in the curved IM. (D, E) A cross-section view and top view of the local refined collar structure in the Δ*motB* motor, respectively. (G, H) Central sections of the asymmetric reconstruction of the WT motor. (I) A section from the unrolled map of the WT motor showing the collar and stator inserted in the curved IM. (J, K) A cross-section view and top view of the local refined collar structure in the WT motor, respectively. (F, L) The FSC curves corresponding to the local refined structures shown in C and H, respectively.

**Figure S5.**
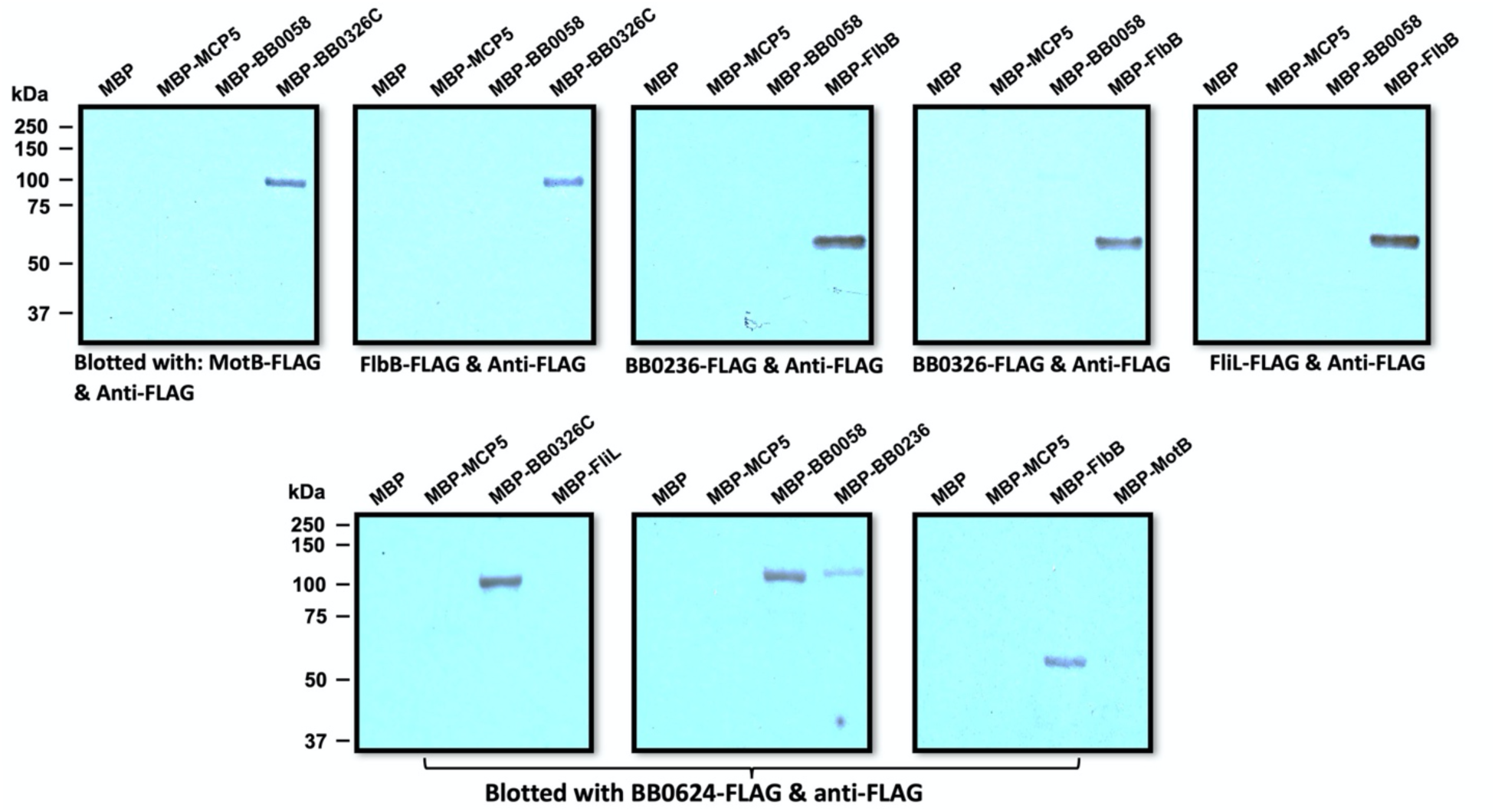
Specific interactions of FlcB (BB0058, top) and FlcC (BB0624, bottom) with other flagellar proteins determined by far western or affinity blotting. Approximately 1 μg of MBP-tagged proteins shown on top of each panel was subjected to SDS-PAGE followed by Coomassie blue staining (not shown) or transferred to a PVDF membrane. The membranes were incubated with 1xFLAG tagged proteins indicated at the bottom of each panel and then immunoblotted with anti-FLAG monoclonal antibodies. FlcB (BB0058) interacts with FlcC; FlcA (BB0326) interacts with MotB; FlcC (BB0624) specifically interacts with FlcA (BB0326), FlcB (BB0058), and MotB.

**Movie S1. The collar changes its conformation to accommodate membrane curvatures in the Δ*motB* motor**. Local refinement and 3D classification (based on the density of the IM) were applied to the Δ*motB* motors. The class averages of the collar region showing the different IM curvatures and conformational changes of the collar subunits. This movie shows four representative class averages with different IM curvatures, highlighting the conformational change of the collar subunit to accommodate diverse membrane curvatures in *B. burgdorferi* cells.

**Movie S2**. Visualization of the modular collar structure and its capacity to accommodate diverse membrane curvatures in *B. burgdorferi* cells.

